# Hydrogel crosslinking mechanisms influence the release and functional delivery of lipid nanoparticles

**DOI:** 10.64898/2026.08.13.741169

**Authors:** Andreas G. Schreiber, Felix Hauswirth, Leon Reger, Simone P. Carneiro, Oliva M. Merkel, Miriam Breunig

**Affiliations:** Department of Pharmaceutical Technology, University of Regensburg, 93053 Regensburg, Germany; Department of Pharmacy, Ludwig-Maximilians-Universität Munich, 81377 Munich, Germany; Institute of Pharmacy and Molecular Biotechnology, Department of Pharmaceutical Technology and Biopharmacy, Heidelberg University, 69120 Heidelberg, Germany

**Keywords:** mRNA-LNPs, hydrogel, prolonged delivery, vaccine delivery

## Abstract

Hydrogels have emerged as attractive vaccine delivery platforms because they enable controlled modulation of antigen availability. However, how different hydrogel environments affect the release and functionality of mRNA-loaded lipid nanoparticles (mRNA-LNPs) remains poorly understood. Here, we investigated the release, stability, cellular uptake, and transfection capability of LNPs released from four hydrogel systems representing distinct crosslinking mechanisms: covalently crosslinked poly(ethylene glycol) (PEG), ionically crosslinked alginate, thermoresponsive Poloxamer 407 (P407), and protein-based Matrigel/collagen hydrogels.

All hydrogel types enabled release of LNPs over days, with lower polymer concentrations generally exhibiting faster nanoparticle release. PEG offered greatest tunability of release kinetics; at the same time the recovery of the LNP-incorporated fluorescent dye DiI was reduced to about 80 %, indicating partial dye leakage. Alginate hydrogels exhibited recovery of DiI below 50 % and broader particle size distributions after release, while P407 hydrogels largely preserved LNP characteristics. CyroTEM analysis revealed that mRNA-LNPs kept at 37 °C, including those released from hydrogels, exhibited a more compact and electron-dense morphology than mRNA-LNPs maintained at 4 °C. Notably, although quantitative recovery from Matrigel/collagen hydrogels was limited, released LNPs achieved approximately 80-90 % of the eGFP expression compared to mRNA-LNPs that have not been embedded. Importantly, cellular uptake and transfection experiments demonstrated that all investigated hydrogels released biologically active mRNA-LNPs capable of mediating protein expression. Moreover, our findings show that hydrogel composition is a critical determinant of mRNA-LNP release, stability, functional delivery and morphology of the released particles. This work provides design principles for the development of hydrogel-based mRNA delivery systems aimed at sustained antigen availability and prolonged vaccine responses.

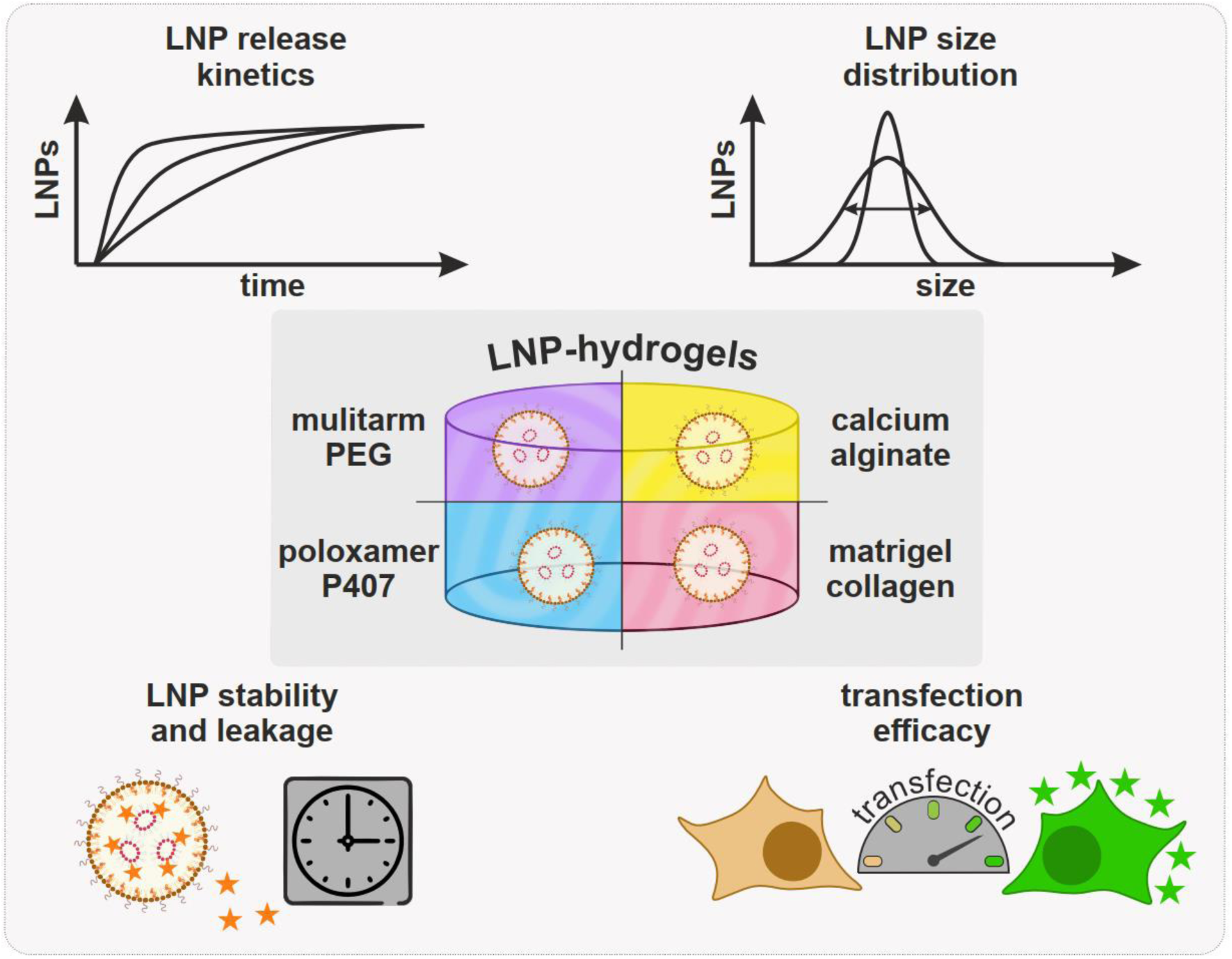

## 1. Introduction

Vaccination has profoundly reduced the global burden of infectious diseases and is increasingly being explored as a therapeutic strategy in cancer immunotherapy [1,2]. The rapid development and worldwide rollout of mRNA vaccines during the COVID-19 pandemic have highlighted the translational potential of nucleic acid-based therapies [3,4]. Lipid nanoparticles (LNPs) play a pivotal role by shielding mRNA from degradation and facilitating its delivery into cells. A major advantage of ionizable lipids in LNP formulations is their ability to promote endosomal escape, thereby enabling cytosolic mRNA delivery and subsequent antigen production [4]. Despite these advances, the transient nature of mRNA expression remains a major limitation for achieving durable antigen presentation and robust immune responses.

Mounting evidence indicates that antigen duration and kinetics are key determinants of vaccine efficacy, critically shaping the magnitude and quality of immune responses [5,6]. Extended antigen exposure enhances germinal center formation, promotes B cell affinity maturation, and improves both the quality and durability of antibody responses [7–9]. Conventional vaccination strategies are typically administered as bolus injections, resulting in a rapid spike in antigen levels followed by swift clearance. Although mRNA-LNP vaccines partly overcome this limitation through transient in situ expression, antigen availability remains relatively short-lived [5,10,11]. In contrast, natural infections provide sustained antigen exposure over extended periods, often accompanied by persistent inflammatory cues [12]. While conventional vaccination strategies are highly effective against many infectious diseases, they frequently fail to induce durable immunity against poorly immunogenic targets including HIV and cancer-associated antigens [8,13,14]. These observations highlight the potential of delivery strategies for sustained antigen presentation.

Various approaches have been explored to prolong antigen availability, including implantable devices [15], microparticle-based systems [16], and microneedle patches [17]. Among these, hydrogels have emerged as particularly attractive platforms due to their injectability, biocompatibility and ability to form local depots for controlled payload release [18–21]. Beyond functioning as delivery reservoirs, hydrogels can actively influence immune responses by recruiting immune cells and establishing localized immunological niches that enhance antigen presentation [22,23]. To date, however, most hydrogel-based vaccine strategies have focused on soluble proteins or protein-based constructs [24–26]. mRNA-LNP delivery from hydrogels remains largely unexplored. To our knowledge, only two studies have investigated such systems in immunological settings, reporting enhanced immune responses associated with prolonged antigen expression and immune cell recruitment [22,27]. These studies primarily report biological outcomes and provide little insight into how hydrogel encapsulation affects mRNA-LNP behavior. In particular, the impact of hydrogel environment on LNP integrity remains poorly characterized, including potential LNP aggregation within the hydrogel or cargo leakage. These interactions are likely to determine not only LNP stability, but also their spatial distribution, transport, and release behavior.

An additional and largely unexplored factor is the role of hydrogel crosslinking chemistry in shaping LNP behavior. Different crosslinking mechanisms generate hydrogel networks that vary widely in stability, mesh architecture, degradability, and molecular interactions. These properties are expected to directly govern the retention and release of encapsulated mRNA-LNPs [28–30], yet their influence has not been systematically investigated. For example, poly(ethylene glycol)- (PEG)-based hydrogels are typically covalently crosslinked, yielding stable and tunable networks with controlled hydrolytic degradation [20,31]. In contrast, Poloxamer 407 (P407) forms physically assembled micellar networks at physiological temperature that are highly dynamic and prone to rapid dissolution [32]. Alginate hydrogels rely on ionic crosslinking through Ca²⁺ mediated junction zones, with network dissolution governed by ion exchange processes [33]. Biomimetic systems such as mixtures of Matrigel/collagen assemble through covalent and non-covalent protein-protein interactions, resulting in heterogeneous and biologically responsive matrices [34].

Here, we investigate how distinct hydrogel crosslinking mechanisms affect the stability, release and functionality of dye- and mRNA-loaded LNPs. We hypothesize that hydrogel network architecture and matrix properties govern LNP transport and retention, thereby controlling release kinetics and biological activity. Because the investigated hydrogels undergo degradation and cargo release within a few days, the focus of this study was not long-term depot formation but rather the modulation of early mRNA-LNP availability. Given the importance of antigen kinetics during the initial phase after vaccination, even short-term prolongation of nanoparticle retention may be sufficient to enhance antigen expression and shape immune priming.

## 2. Materials and methods

### 2.1. Materials

1,2-Distearoyl-sn-glycero-3-phosphocholine (DSPC) was kindly provided by Lipoid GmbH (Ludwigshafen, Germany). Cholesterol was purchased from Carl Roth GmbH (Karlsruhe, Germany). ALC-0315 and ALC-0159 were obtained from Biomol GmbH (Hamburg, Germany). Acetic acid (glacial, p.a.), ethanol (p.a.), trypsin-EDTA, 4’,6-diamidino-2-phenylindole (DAPI), paraformaldehyde (PFA), Poloxamer 407 (P407), ethidium bromide, propidium iodide and Triton X-100 were purchased from Sigma-Aldrich (Merck KGaA, Darmstadt, Germany). Phosphate-buffered saline (PBS), Dulbecco’s Modified Eagle Medium (DMEM), and penicillin-streptomycin solution were obtained from Gibco (Thermo Fisher Scientific, Waltham, MA, USA). Fetal bovine serum (FBS) was purchased from PAN-Biotech GmbH (Aidenbach, Germany). Dako Fluorescence Mounting Medium was obtained from Agilent Technologies (Santa Clara, CA, USA). Sodium alginate (Protanal LF 10/60 FT) was supplied by FMC Biopolymer (Philadelphia, PA, USA). Collagen and Matrigel Basement Membrane Matrix were purchased from Corning Inc. (Corning, NY, USA). Agarose was obtained from Biozym Scientific GmbH (Hessisch Oldendorf, Germany). DiI was obtained from Lumiprobe GmbH (Hannover, Germany). eGFP-encoding, 5’ Cap1 mRNA was purchased from Ribopro GmbH (Leeuwarden, The Netherlands). Diels-Alder crosslinkable 4armPEG_40k_-maleimide and 4armPEG_40k_-furan derivatives were synthesized as previously described in [20]. The degree of functionalization, determined by ^1^H NMR spectroscopy, were 84 % for both, 4armPEG_40k_-maleimide and 4armPEG_40k_-furan. All aqueous solutions were prepared using Milli-Q water.

### 2.2. LNP formulation

LNPs were generated by microfluidic mixing using a NanoAssemblr Ignite system (Precision Nanosystems Inc., Vancouver, Canada, now part of Cytiva, Massachusetts, USA). Lipid components were dissolved in ethanol at a total lipid concentration of 5 mM at a molar ratio of DSPC:cholesterol:ALC-0315:ALC-0159 = 9.4:42.7:46.3:1.6, similar to the composition of the Comirnaty (BioNTech/Pfizer, Mainz, Germany) formulation. The organic (ethanolic) lipid phase and the aqueous phase (25 mM acetate buffer, pH = 4) were combined at a flow rate ratio of 3:1 (aqueous:organic) and a total flow rate of 12 mL/min. For dye-labeled formulations, DiI was co-dissolved in the lipid phase at 0.8 mol%. Only for the DiI concentration series used for Stern-Volmer analysis, LNPs were generated using a Sunscreen microfluidic mixing system (Unchained Labs, Pleasanton, CA, USA) operated at a flow rate ratio of 3:1 (aqueous:organic) and a total flow rate of 8 mL/min. For mRNA-loaded formulations, encapsulation was performed at a nitrogen-to-phosphate (N:P) ratio of 6. Following microfluidic mixing, LNP suspensions were collected and dialyzed against PBS (pH 7.4) overnight on ice to remove ethanol and adjust the buffer conditions.

### 2.3. LNP characterization

Particle size and polydispersity index (PDI) were determined by dynamic light scattering (DLS) using a Zetasizer Nano ZS (Malvern Panalytical, Malvern, UK) at 25 °C. ζ-potential was measured by laser Doppler anemometry on the same instrument after dilution of the samples 1:20 in Milli-Q water. Particle size distribution and concentration were further analyzed by nanoparticle tracking analysis (NTA) using a NanoSight NS300 (Malvern Panalytical, Malvern, UK) at 25 °C. The span of the NTA size distribution was calculated as span = (D_90_-D_10_)/D_50_, where D_10_, D_50_, and D_90_ represent the 10^th^, 50^th^, and 90^th^ percentiles, respectively. Fluorescence self-quenching was determined using DiI-LNP formulations diluted either with PBS for fluorescence of intact DiI-LNPs or 2 % (v/v) Triton X-100 for total fluorescence in a 384-well plate. After incubation for 10 min, fluorescence intensity was measured using a BioTek SYNERGY neo2 fluorescence reader (Agilent Technologies, Santa Clara, USA). Total mRNA recovery and encapsulation efficiency of mRNA-LNPs were determined using the Quant-iT RiboGreen assay (Invitrogen, Thermo Fisher Scientific, Schwerte, Germany) in a 384-well plate. mRNA-LNPs were diluted with either TE buffer for the determination of encapsulation efficiency or 2 % (v/v) Triton X-100 in TE buffer for total mRNA recovery. Subsequently, an equal volume of RiboGreen reagent diluted 1:200 was added to each well. After incubation for 10 min, fluorescence intensity was measured using a FLUOstar Omega fluorescence reader (BMG LABTECH, Ortenberg, Germany) using 480/10 nm excitation and 520/10 nm emission filters. Encapsulation efficiency (EE) was calculated using following formula:

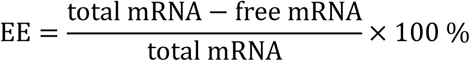

Agarose gels (1 % w/v) were prepared with 1× TAE buffer and 0.8 µg/mL ethidium bromide. For analysis of encapsulated mRNA, mRNA-LNPs were lysed using a TE buffer containing 2 % (v/v) Triton X-100, analogous to the procedure employed for the RiboGreen assay. Samples were adjusted to an mRNA concentration of approximately 8 µg/mL. Prior to electrophoresis, samples were supplemented with 10× loading buffer to yield a final loading buffer concentration of 1×. Approximately 0.1 µg mRNA were loaded per well. Electrophoresis was performed in 1× TAE running buffer, and nucleic acids were visualized by UV illumination using the incorporated ethidium bromide stain.

### 2.4. Hydrogel preparation

Hydrogels were prepared in a total volume of 50 µL containing 15 µL of either DiI-LNPs or mRNA-LNPs (equivalent to ≈ 500 ng mRNA per Gel). For PEG-based hydrogels, Diels-Alder crosslinkable PEG derivatives were dissolved at 30 % (w/w) in 50 mM phosphate buffer (pH 5) to minimize hydrolysis of the maleimide functional groups. Alginate hydrogels were prepared from a 4 % (w/w) stock solution in Milli-Q water and mixed with LNPs. For gelation, 45 µL of the alginate-LNP mixture was quickly added to 5 µL of 100 mM CaCl_2_. Poloxamer 407 (P407) hydrogels were prepared by dissolving the polymer at 45 % (w/w) in Milli-Q water on ice. For Matrigel/collagen hydrogels, collagen solutions were first neutralized using 0.1 N NaOH and subsequently mixed with Matrigel (final concentration in gel 2 mg/mL) on ice. While in case of the PEG hydrogels, gelation was allowed to proceed overnight at 37 °C. All other hydrogel mixtures were formed by incubation at 37 °C for at least 30 min.

### 2.5. In vitro release study

Gelation was performed in 1.5 mL microtubes. Following gelation, 350 µL of pre-warmed PBS was added to each sample to achieve a final volume of 400 µL. Samples were incubated at 37 °C under gentle agitation (50 rpm). At defined time points, 20 µL aliquots were withdrawn and replaced with an equal volume of fresh PBS to maintain sink conditions. Collected samples were analyzed for fluorescence on a BioTek SYNERGY neo2 fluorescence reader (Agilent Technologies, Santa Clara, USA) using 543/15 nm excitation and 580/15 nm emission filters.

### 2.6. cryoTEM characterization

Prior to imaging, reference LNPs and hydrogel-released LNPs were concentrated using Amicon Ultra-4 Ultracel 100 kDa centrifugal filter units. For reference LNPs incubated at 37 °C and hydrogel-released samples, 400 μL of hydrogel supernatant was subjected to centrifugal filtration at 2000 rcf for 20 min, resulting in an approximately 5 to 10-fold concentration of the nanoparticles. As described in [35], 3.5 μL of each concentrated sample was deposited onto 2 nm carbon-coated Quantifoil R3/3 copper grids and vitrified using a Vitrobot Mark IV (Thermo Fisher Scientific). CryoTEM images were acquired using a Tecnai G2 Spirit transmission electron microscope (Thermo Fisher Scientific) operated at 120 kV and equipped with a 2048 × 2048 pixel CCD camera (TVIPS GmbH). Image acquisition was performed using the EM-TOOLS software package (TVIPS GmbH).

### 2.7. In vitro LNP cell uptake studies from hydrogel release

HeLa cells (human cervical carcinoma cell line) were cultured in DMEM supplemented with 10 % fetal bovine serum (FBS) and 1 % penicillin-streptomycin. Cells were maintained at 37 °C in a humidified incubator with 5 % CO₂. Upon reaching ∼ 80 % confluency, cells were passaged using standard trypsinization procedures (trypsin-EDTA).

For epifluorescence microscopy, HeLa cells were seeded in glass-bottom 24-well plates at a density of 7.5 × 10^3^ cells per well in 500 µL of DMEM supplemented with 10 % FBS and cultured for 24 h to allow cell attachment. Cellular uptake of LNPs released from hydrogels was investigated using 6.5 mm Transwell inserts with a 5.0 µm pore polycarbonate membrane (Costar #3421, Corning, Corning, NY, USA). LNP-loaded hydrogel sols were prepared as described above and cast (50 µL) into the Transwell inserts. After gelation, the gels were overlaid with 100 µL culture medium and transferred into wells containing the pre-seeded cells. Co-incubation was performed for 72 h at 37 °C to enable release of LNPs from the hydrogels and subsequent cellular uptake. Following incubation, the Transwell inserts were removed, and all subsequent steps were performed with the cells remaining in the glass-bottom well plates. Cells were washed with PBS, fixed with 4 % paraformaldehyde for 15 min at room temperature, and stained with DAPI for nuclear visualization. Following washing, cells were mounted and stored protected from light until imaging. Fluorescence images were acquired using a Zeiss Axio Observer 7 epifluorescence microscope equipped with a 63× objective (Carl Zeiss, Jena, Germany).

To quantify functional mRNA delivery following exposure to collected hydrogel degradation supernatants, HeLa cells were seeded in 24-well plates at a density of 50 × 10^3^ cells per well in 500 µL of DMEM supplemented with 10 % FBS and cultured for 24 h prior to transfection. mRNA-LNP-loaded hydrogels were prepared and incubated under the same release conditions as described for the in vitro release study. Following 72 h of release at 37 °C, 100 µL of the collected degradation supernatant were added to HeLa cells cultured in 24-well plates containing 400 µL of fresh culture medium per well. Cells were incubated for an additional 24 h at 37 °C to allow eGFP expression. Subsequently, cells were washed with PBS, detached by trypsinization, and resuspended in culture medium to inactivate trypsin. Cell suspensions were transferred to reaction tubes and centrifuged at 200 rcf for 5 min at 4 °C. The cell pellets were washed once with cold PBS and finally resuspended in 300 µL PBS. Samples were kept on ice until analysis on a FACS Canto II flow cytometer (BD Biosciences, Franklin Lakes, NJ, USA). For the assessment of hydrogel-associated cytotoxicity, hydrogels without LNPs were incubated under release conditions as described above. Following resuspension in PBS, 20 µL propidium iodide solution (10 µg/mL) was added, and samples were incubated for 1 min prior to analysis.

To evaluate functional mRNA delivery under conditions of continuous hydrogel-mediated nanoparticle release, mRNA-LNP-loaded hydrogels were incubated in Transwell inserts as described above for the cellular uptake experiments. HeLa cells were seeded at a density of 7.5 × 10³ cells per well. As a reference, non-encapsulated mRNA-LNPs were added directly to the insert compartment above the membrane. Following 72 h of co-incubation, cells were harvested and analyzed by flow cytometry as described above.

### 2.8. Statistical analysis

Results are expressed as mean ± standard deviation of triplicate experiments. Statistical analyses and curve fitting were performed using Origin 2023b software (OriginLab Corporation, Northampton, MA, USA).

## 3. Results

### 3.1. Physicochemical LNP characteristics

To investigate interactions between LNPs and hydrogel matrices, we employed dye-loaded particles as a surrogate system alongside mRNA-loaded formulations. This approach enables direct probing of LNP behavior independent of cargo-related biological effects.

As a first step, we determined the optimal loading conditions for the lipophilic fluorescent probe DiI. LNPs were prepared with dye concentrations ranging from 0.1 to 1 mol% to identify maximal dye-loading with minimal self-quenching. Total fluorescence obtained after particle lysis, F_0_, was normalized to the fluorescence of intact LNPs, F, according to the classical Stern-Volmer relationship [36]. Under the self-quenching regime applied here, the incorporated dye itself acts as quencher, such that the ratio F_0_/F reflects the extent of fluorescence quenching at increasing dye content. As shown in Fig. 1A, fluorescence quenching increased approximately linearly with dye loading. The corresponding Stern-Volmer constant (K_SV_ ≈ 1 mol%^−1^) indicates a modest but detectable self-quenching efficiency. To monitor LNPs at the single-particle level and to determine LNP concentration after release from hydrogels, nanoparticle tracking analysis (NTA) was performed in the fluorescence mode. The method has the great advantage that it allows to detect fluorescently labeled LNPs in the presence of a noisy background, e.g. emerging from hydrogel degradation products. The signal-to-noise ratio (S/N) of NTA measurements decreased sharply with decreasing dye content, and reliable particle discrimination from background was no longer possible below 0.6 mol% DiI. Based on these results, a dye loading of 0.8 mol% was selected for subsequent experiments. Here, particle concentrations determined by NTA in fluorescence mode were consistent with those obtained in conventional light-scattering mode (Fig. S1), both yielding values of approximately 1 × 10^12^ LNPs/mL. Following validation of the DiI-LNP formulation, the corresponding mRNA-loaded LNPs were characterized with respect to RNA recovery and encapsulation. The applied formulation protocol achieved complete mRNA recovery and an encapsulation efficiency of 82.0 (±1.8) % (Fig. 1B). Furthermore, agarose gel electrophoresis revealed no discernible differences between free mRNA and mRNA recovered from mRNA-LNPs (Fig. S2). Physicochemical characterization revealed comparable size distributions for dye-loaded and mRNA-loaded LNPs, with hydrodynamic diameters of about 50 nm and low polydispersity (below 0.15) (Fig. 1C). ζ-potential measurements indicated moderately negative surface charges for both formulations, with mRNA-LNPs displaying a slightly more pronounced negative potential compared to their dye-loaded counterparts (Fig. 1D).

**Fig. 1.**
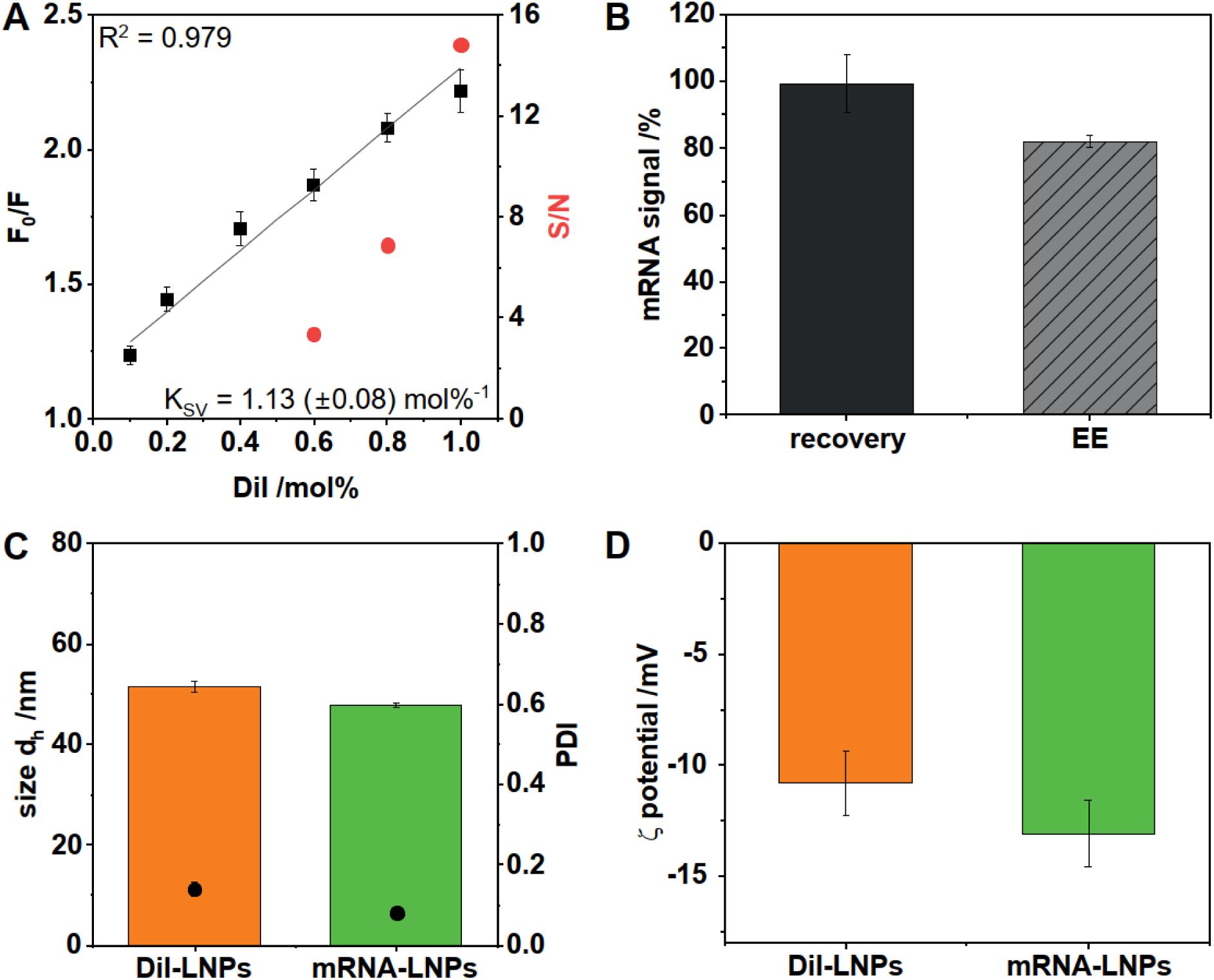
Physicochemical characterization of DiI- and mRNA-loaded LNPs. A) Self-quenching analysis of DiI fluorescence at increasing dye loading. The ratio of total fluorescence after lysis, F_0_, to fluorescence of intact particles, F, follows a Stern-Volmer relationship, showing an approximately linear increase in quenching with dye concentration, indicative of moderate non-radiative relaxation. Nanoparticle tracking analysis (NTA) in fluorescence mode reveals a strong dependence of signal-to-noise ratio (S/N) on dye loading, with optimal detectability starting from 0.8 mol%. B) The initially applied mRNA was completely recovered in the final mRNA-LNP formulation, with an encapsulation efficiency exceeding 80 % C) Size distribution analysis by DLS shows hydrodynamic diameters around 50 nm with low polydispersity for both dye- and mRNA-loaded LNPs, with mRNA-containing particles exhibiting slightly larger size. D) ζ-potential measurements indicate moderately negative surface charges, slightly more pronounced for mRNA-LNPs. Data are presented as mean ± SD.

### 3.2. In vitro release kinetics of DiI-LNPs

To investigate the release behavior of LNPs from hydrogel depots, DiI-labeled particles were incorporated into the four hydrogel systems with distinct crosslinking mechanisms: covalently crosslinked 4armPEG_40k_ (from now on only dedicated as PEG) hydrogels, ionically crosslinked alginate gels, thermoresponsive Poloxamer 407 (P407), and protein-based Matrigel/collagen matrices. Release studies were performed at 37 °C under gentle agitation. At the same time, the stability of DiI-LNPs was assessed under conditions matching the release experiments using reference LNPs, which throughout this study are defined as LNPs handled at the same temperature (37 °C) and concentration as the corresponding experimental samples. This reference is particularly important as mRNA encapsulated within LNPs can undergo degradation even during storage at 4 °C [37,38], making exposure to identical incubation conditions essential for distinguishing temperature-induced effect from hydrogel-related changes. Both fluorescence intensity (Fig. S3A) and particle concentration determined by fluorescence NTA (Fig. S3B) remained constant throughout the duration of the experiment, indicating that the DiI-LNPs remained stable and no dye leakage occurred at incubation conditions.

As shown in Fig. 2, all hydrogel systems exhibited concentration-dependent release profiles of the LNP-incorporated dye DiI over approximately three days. Time points at which complete macroscopic gel degradation was observed are indicated by asterisks. Across all materials, lower polymer concentrations accelerated the release of DiI-associated fluorescence, although pronounced differences were observed between hydrogel classes. Among the investigated systems, covalently crosslinked PEG hydrogels exhibited the most tunable release profiles and yielded a recovery of up to approximately 80 % of the initially incorporated fluorescent dye (Fig. 2A). In contrast, alginate hydrogels yielded substantially lower fluorescence recoveries, with less than 50 % of the initially incorporated signal detected after complete macroscopic gel degradation (Fig. 2B). P407 hydrogels showed the strongest concentration dependence, with recoveries of DiI ranging from approximately 60 % to a nearly quantitative amount (Fig. 2C). Notably, rapid dissolution of the 20 % P407 formulation was accompanied by the highest fluorescence recovery observed across all hydrogel systems. The Matrigel/collagen system demonstrated a substantial loss of the DiI fluorescence (Fig. 2D). As complete matrix dissolution was not achieved under the experimental conditions, released LNPs could not be completely separated from residual hydrogel material, likely contributing to an underestimation of fluorescence recovery. Importantly, the assay quantified the fluorescence signal associated with DiI-labeled LNPs rather than intact LNPs. Consequently, incomplete recovery of the fluorescence signal does not necessarily indicate nanoparticle loss but may also result from partial leakage of DiI from the LNPs.

**Fig. 2.**
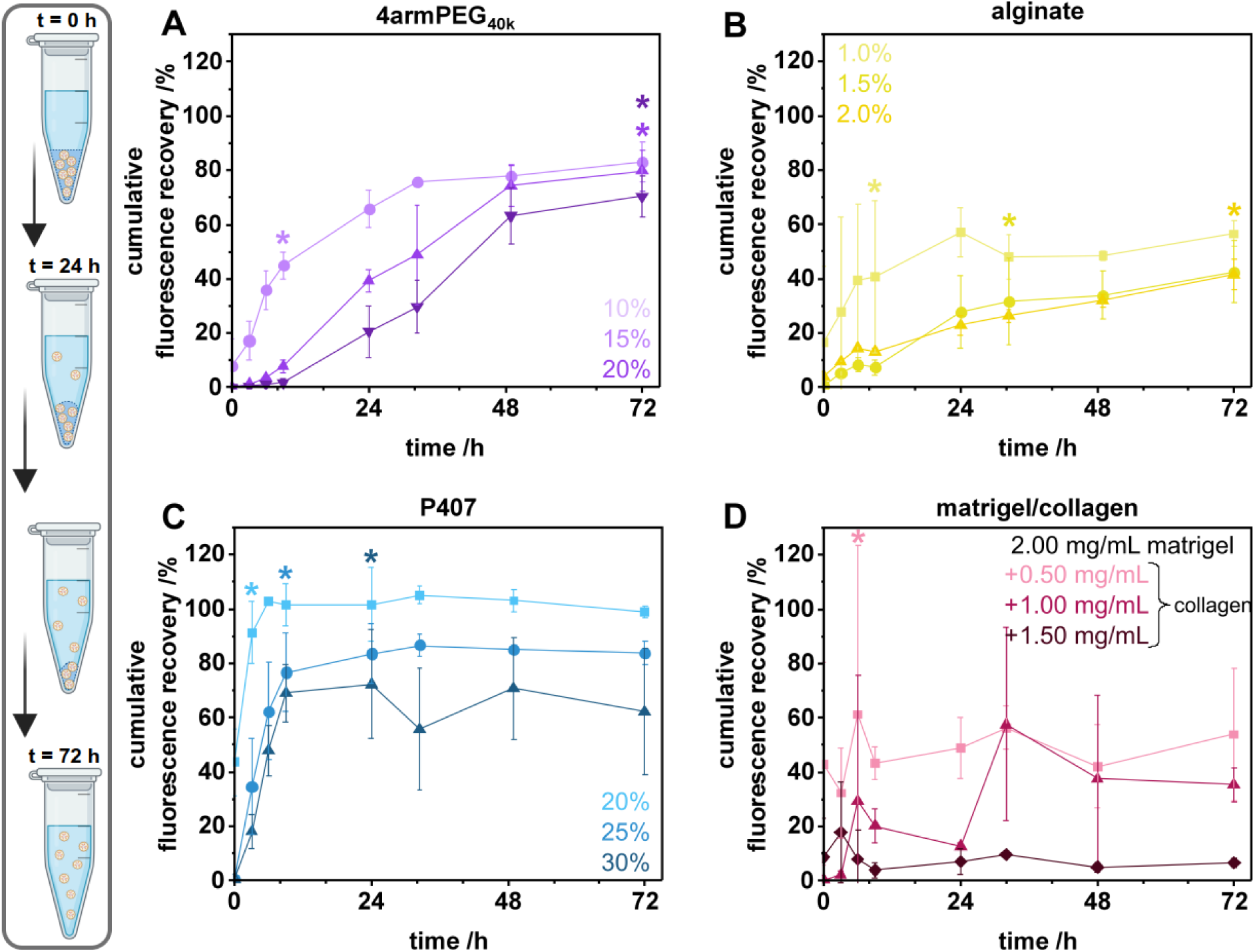
Hydrogel-dependent in vitro release kinetics. DiI-LNPs were incorporated into four hydrogel platforms with distinct crosslinking mechanisms: covalently crosslinked 4armPEG_40k_, ionically crosslinked alginate, thermoresponsive P407, and protein-based Matrigel/collagen hydrogels. Release was monitored at 37 °C under gentle agitation. All hydrogel systems exhibited concentration-dependent release kinetics, with lower polymer concentrations resulting in faster nanoparticle release. Recovery of released LNPs varied substantially between materials, reaching approximately A) 80 % for PEG hydrogels, B) 50 % for alginate, and C) 60-100 % for P407 depending on polymer concentration. D) Matrigel/collagen hydrogels showed incomplete fluorescence recovery due to incomplete matrix dissolution under the experimental conditions. Asterisks indicate time points at which complete macroscopic gel degradation was observed by visual inspection. Created in BioRender. Breunig, M. (2026) https://BioRender.com/ccgcuvf. Data are presented as mean ± SD.

### 3.3. Effect of hydrogel encapsulation on LNP size, morphology and particle recovery

Subsequent NTA analysis was therefore performed to determine whether reduced fluorescence recovery was associated with particle loss or merely reflected partial dye leakage while cryoTEM imaging was used to assess potential changes in particle morphology. Again, as a comparison, reference LNPs were incubated at 37 °C for the duration of the release experiments to account for potential temperature induced changes independent of the hydrogel environment.

NTA revealed hydrogel- and concentration-dependent changes in the particle size distributions (Fig. 3A). The reference formulation exhibited a modal hydrodynamic diameter of 62 (±3) nm and a span value below 1, indicating a narrow and homogeneous particle size distribution. Across all hydrogel systems, the modal particle diameter remained largely unchanged, indicating that encapsulation did not induce substantial particle growth or aggregation. For PEG, increasing polymer concentration shifted the mode towards smaller particle sizes and increased the span. Alginate showed a slight concentration-dependent increase in modal size, accompanied by consistently elevated span values. P407 resulted in slightly smaller modal sizes than the reference LNPs, while the span increased with polymer concentration. Particle concentrations after release were determined by fluorescence NTA (Fig. 3B), enabling direct quantification of DiI-labeled nanoparticles. Across all hydrogel formulations, concentration recoveries ranged between approximately 100 and 150 %. The findings indicate that hydrogel encapsulation did not result in substantial particle loss. Values above 100 % are within the expected variability of NTA measurements [37]. Fluorescence NTA nevertheless provides a robust estimate of nanoparticle recovery because only DiI-labeled particles are detected, thereby reducing interference from hydrogel-derived debris and other background material. Consequently, the incomplete recoveries observed in the fluorescence-based release experiments (Fig. 2) likely reflect, at least in part, leakage of DiI rather than loss of intact nanoparticles. Overall, hydrogel encapsulation had little effect on average particle size or nanoparticle recovery but increased particle heterogeneity in a hydrogel-dependent manner.

**Fig. 3.**
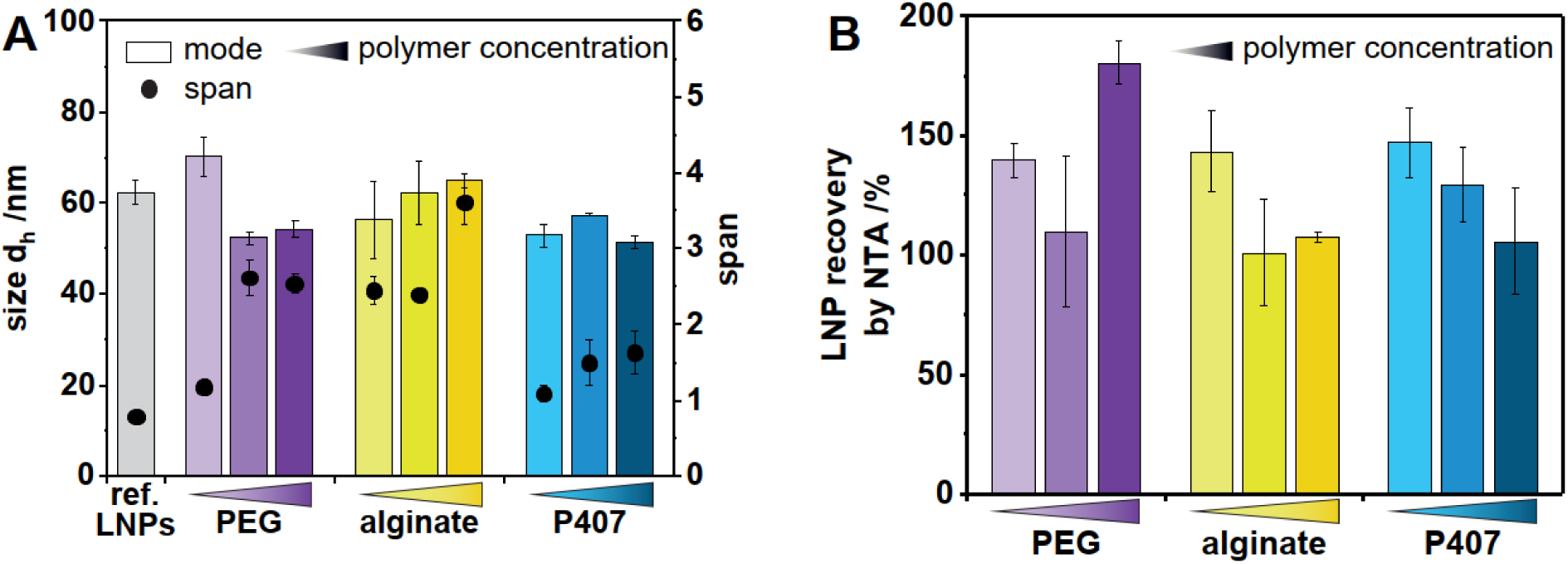
Effect of hydrogel encapsulation on the size distribution and particle recovery of released DiI-LNPs determined by fluorescence NTA. A) While the modal particle diameter remained largely unchanged across all formulations, released DiI-LNPs generally exhibited broader size distributions, reflected by increased span values. B) Particle recoveries ranged between approximately 100 and 150 % for all analyzed hydrogels, indicating efficient particle release and minimal particle loss. Data are presented as mean ± SD.

CryoTEM micrographs revealed a clear difference between particles stored at 4 °C (Fig. 4A) or having been incubated at 37 °C in the absence (reference LNPs, Fig. 4B) or presence of a hydrogel (Fig. 4C-E). LNPs stored at 4 °C exhibited the expected morphology with some particles showing biphasic bleb-like structures [39,40]. mRNA-LNPs incubated at physiological temperature, including hydrogel-released particles and reference LNPs, appeared darker and more compact than corresponding LNPs stored state at 4 °C, indicating increased electron density and suggesting structural compaction of the particles. Quantitative analysis of the images confirmed this observation (Fig. 4F). LNPs stored at 4 °C exhibited larger diameters than reference LNPs incubated at 37 °C as well as LNPs released from the hydrogels. Thus, cryoTEM demonstrated a reduction in particle diameter following incubation at physiological temperature even in the absence of hydrogel exposure.

**Fig. 4.**
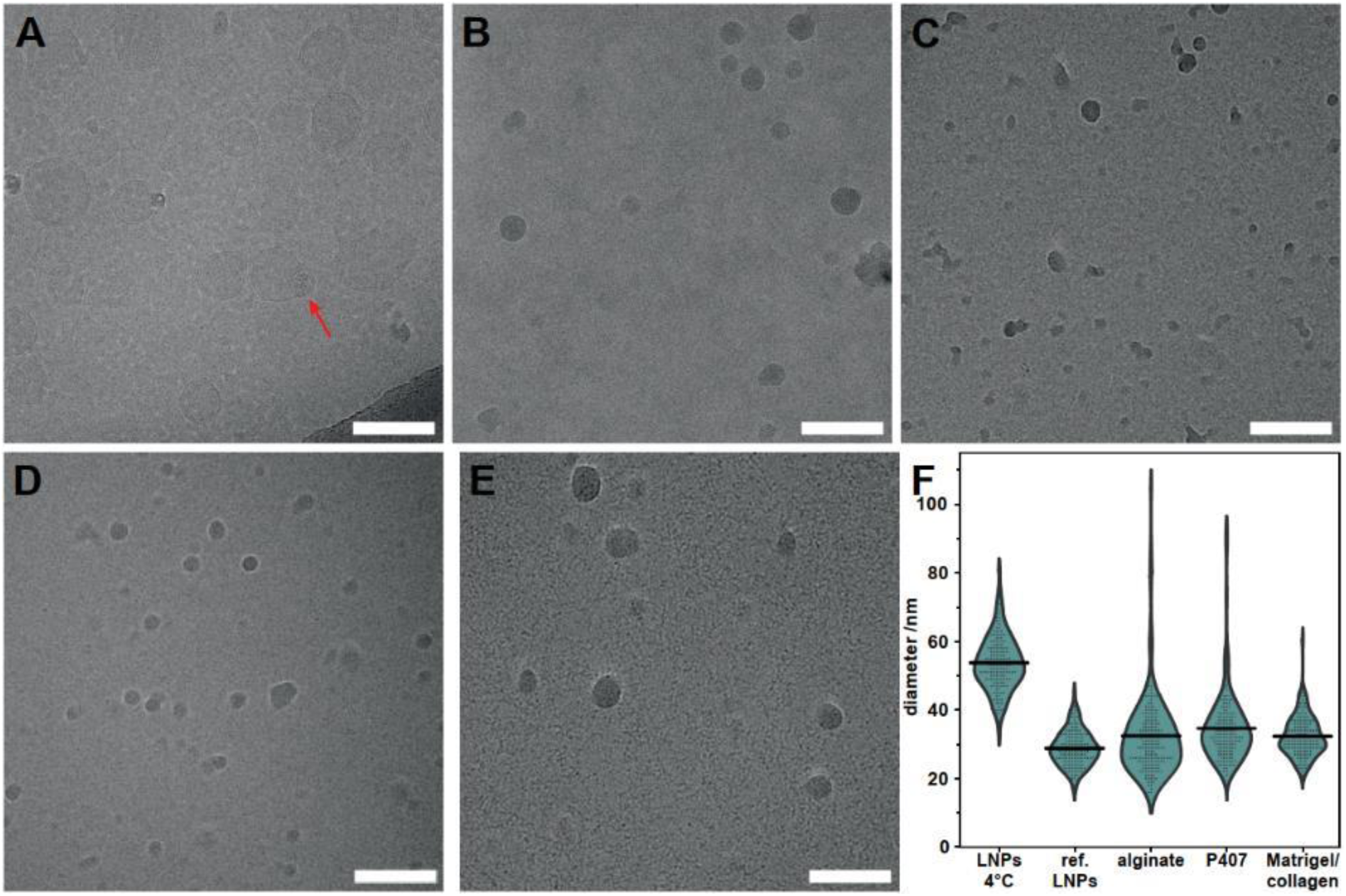
CryoTEM micrographs of mRNA-LNPs. Representative cryoTEM images of A) LNPs from standard storage conditions at 4 °C, B) reference LNPs incubated at 37 °C and LNPs recovered followed release from C) alginate, D) P407 and E) Matrigel/collagen hydrogels. Compared with mRNA-LNPs stored at 4 °C, particles incubated at 37 °C, including reference and hydrogel-released mRNA-LNPs, appeared more compact and electron-dense. The red arrow indicates a bleb-like structure. F) Violin plots confirmed reduced particle diameters for all samples incubated at 37 °C compared with mRNA-LNPs stored at 4 °C. Individual data points represent single LNPs, and black lines indicate mean diameters. PEG hydrogels are not included, as the sample failed to vitrify during cryoTEM sample preparation. Scale bars represent 100 nm.

### 3.4. Particle uptake following continuous release of hydrogel-embedded DiI-LNPs in a Transwell setup

To assess whether hydrogel-released LNPs remained available for cellular uptake, DiI-LNP-loaded hydrogels were placed in Transwell inserts above cultured HeLa cells. This setup allowed released LNPs to diffuse across the membrane and reach the underlying cell layer. As shown in Fig. 5, intracellular DiI fluorescence was detected for all hydrogel systems, confirming that released LNPs remained accessible to cells following release. However, the extent of uptake strongly depended on both hydrogel type and polymer concentration. PEG hydrogels exhibited the most pronounced concentration dependence, with increasing polymer content resulting in progressively lower cellular fluorescence. A similar, albeit weaker, trend was observed for alginate hydrogels, which overall generated the lowest fluorescence signals. Notably, alginate gels did not fully degrade within the Transwell inserts during the experiment. In contrast, P407 and Matrigel/collagen hydrogels led to strong intracellular fluorescence across all concentrations, with no clear concentration-dependent effects. These findings indicate that hydrogel-dependent differences in LNP release translated directly into differences in LNP availability for cellular uptake.

**Fig. 5.**
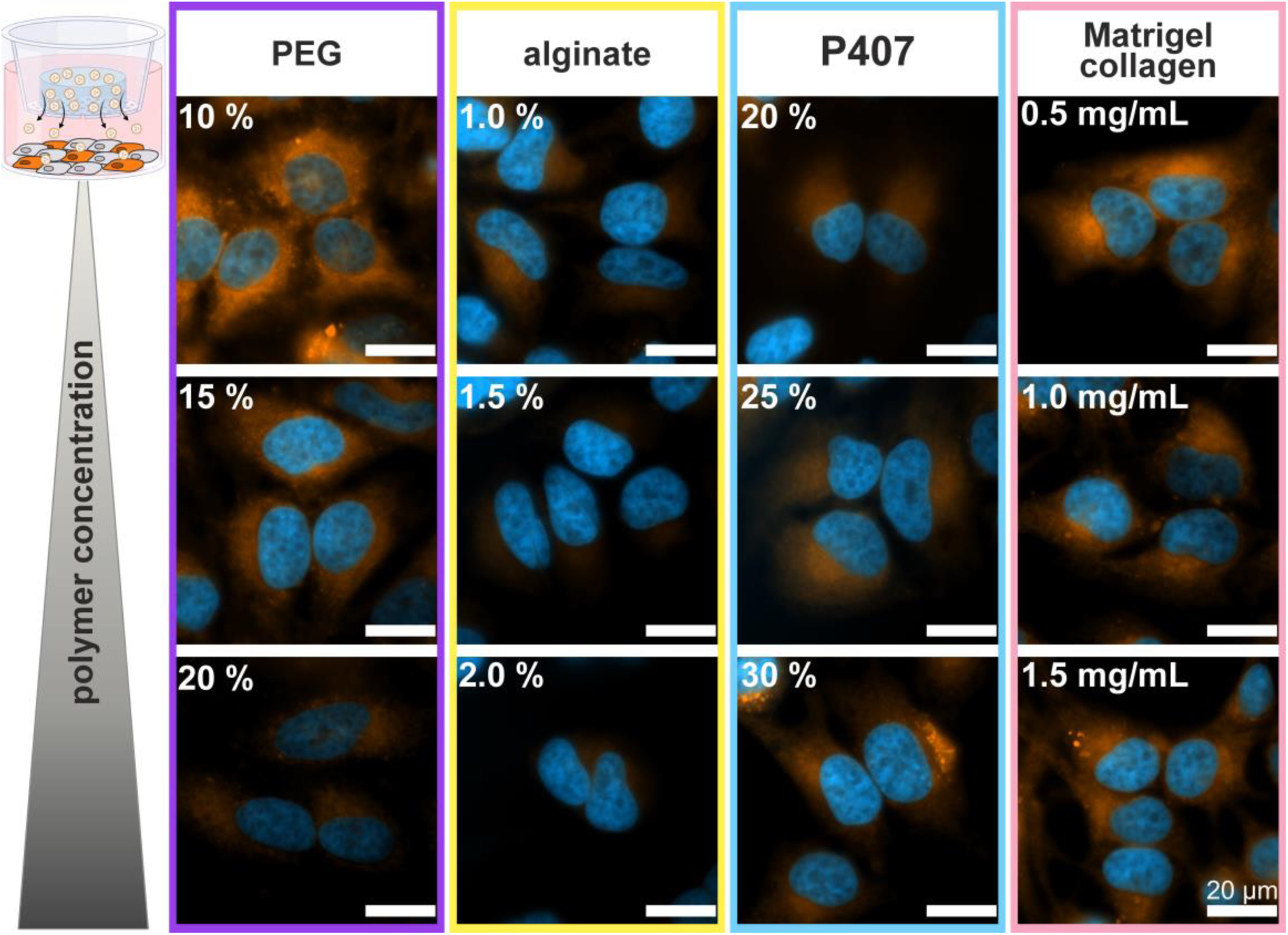
Hydrogel-dependent uptake internalization of released DiI-LNPs in HeLa cells. Hydrogels loaded with DiI-LNPs were cast into Transwell inserts and incubated above HeLa cells for 72 h. Cellular uptake of released nanoparticles was analyzed by epifluorescence microscopy. PEG hydrogels exhibited a strong concentration-dependent decrease in cellular fluorescence, whereas alginate hydrogels yielded overall weaker signals that further declined with increasing polymer concentration. In contrast, P407 and Matrigel/collagen hydrogels produced strong fluorescence signals at all concentrations without a clear concentration-dependent trend. Created in BioRender.com. Breunig, M. (2026) https://biorender.com/dy6uto8.Scale bars represent 20 µm.

### 3.5. Functional transfection by hydrogel-released mRNA-LNPs

#### 3.5.1. Activity of mRNA-LNPs collected from degradation supernatants

To determine whether released mRNA-LNPs retained their biological activity, HeLa cells were exposed to LNPs collected from hydrogel supernatants after 72 h, and eGFP expression was quantified by flow cytometry. eGFP fluorescence intensities were normalized to reference mRNA-LNPs that have been incubated at 37 °C for the duration of the release experiments to account for temperature-induced changes independent of the hydrogel environment. All hydrogel systems enabled functional mRNA delivery, resulting in detectable eGFP expression (Fig. 6A). eGFP expression generally decreased with increasing polymer concentration, indicating that hydrogel composition and network density strongly influenced the availability of functional mRNA-LNPs. Among the investigated formulations, PEG, alginate, and P407 hydrogels yielded comparable expression levels. The Matrigel/collagen interpenetrating system exhibited the highest transfection performance, with the lowest and intermediate collagen supplementation resulting in fluorescence intensities comparable to those observed for the reference formulation. The percentage of eGFP-positive cells, i.e. the transfection rate, showed a similar trend as the eGFP expression and declined with increasing polymer concentration (Fig. 6B). Moreover, PEG hydrogels induced a slight concentration-dependent reduction in cell viability; however, viability remained above 90 % across all fromulations (Fig. S4). Alginate hydrogels similarly caused a modest decrease in viability, although no clear concentration dependence was observed. In contrast, P407 and Matrigel/collagen hydrogels did not affect cell viability at all. Together, these findings demonstrate that all investigated hydrogels released biologically active mRNA-LNPs capable of mediating protein expression without causing substantial cytotoxicity.

**Fig. 6.**
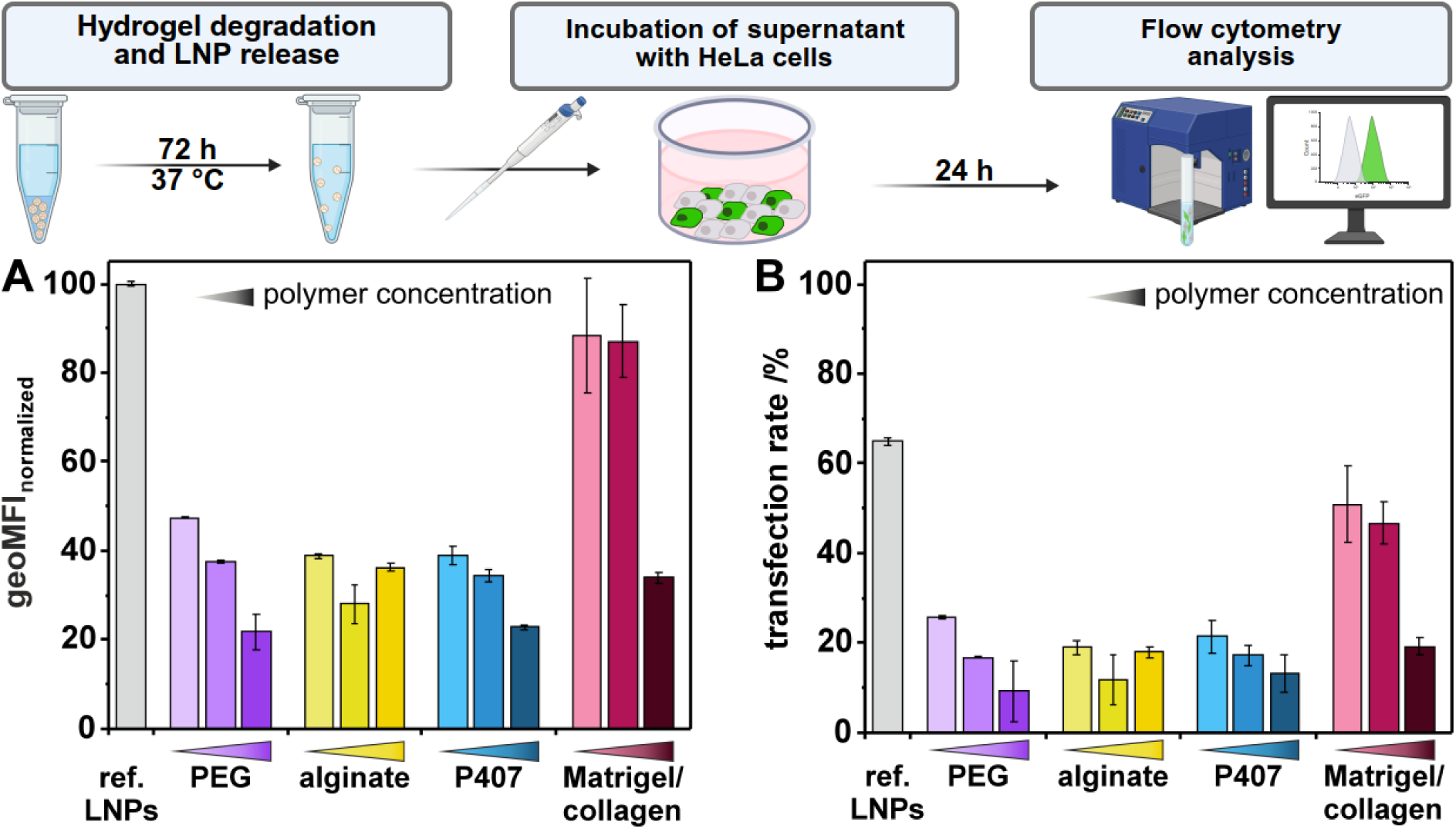
Functional transfection of HeLa cells by hydrogel-released mRNA-LNPs. The supernatants of mRNA-LNP-loaded hydrogels after degradation for 72 h at 37 °C were applied to HeLa cells. After 24 h incubation, eGFP expression was quantified by flow cytometry. mRNA-LNPs that had been incubated under similar conditions at 37 °C served as reference. A) Geo mean fluorescence intensity (geoMFI) normalized to the reference mRNA-LNPs. All hydrogel systems enabled functional mRNA delivery, with eGFP expression generally decreasing at higher polymer concentrations. B) Transfection rates closely mirrored the geoMFI data and declined with increasing polymer concentration. Created in BioRender. Breunig, M. (2026) https://BioRender.com/m4dzfmu. Data are presented as mean ± SD.

#### 3.5.2. Transfection following continuous release of hydrogel-embedded mRNA-LNPs in a Transwell setup

To assess the impact of continuous hydrogel-mediated mRNA-LNP release on cellular transfection, HeLa cells were cultured in the lower compartment of Transwell inserts, while hydrogel formulations containing mRNA-LNPs were placed directly into the inserts. As a reference, non-encapsulated mRNA-LNPs were added directly to the insert compartment above the membrane, allowing comparison with hydrogel formulations under otherwise identical Transwell culture conditions. The hydrogels were incubated for three days, enabling continuous release of mRNA-LNPs and gaining access to the cells via the pores of the Transwell inserts.

Reference mRNA-LNPs resulted in the highest eGFP expression and were defined as 100 % eGFP fluorescence signal. Overall, the formulations followed the order reference LNPs ≈ Matrigel > PEG > alginate > P407 with respect to both eGFP expression (Fig. 7A) and transfection efficiency (Fig. 7B). PEG hydrogels maintained high signal strength, although a moderate reduction compared to the reference was observed. In contrast, alginate hydrogels markedly reduced fluorescence intensity, whereas P407 hydrogels produced only minimal eGFP expression with values close to background levels. Matrigel/collagen hydrogels preserved reporter expression at a level comparable to the reference formulation. A similar trend was observed for the proportion of eGFP-positive cells, i.e. the transfection rate (Fig. 7B). But here, Matrigel/collagen matrices maintained levels of eGFP-positive cells comparable to, or slightly exceeding, those achieved by the reference mRNA-LNPs. Additional control experiments comparing direct mRNA-LNP administration with Transwell-mediated delivery demonstrated that diffusion across the Transwell membrane only minimally affected transfection performance (Fig. S5).

**Fig. 7.**
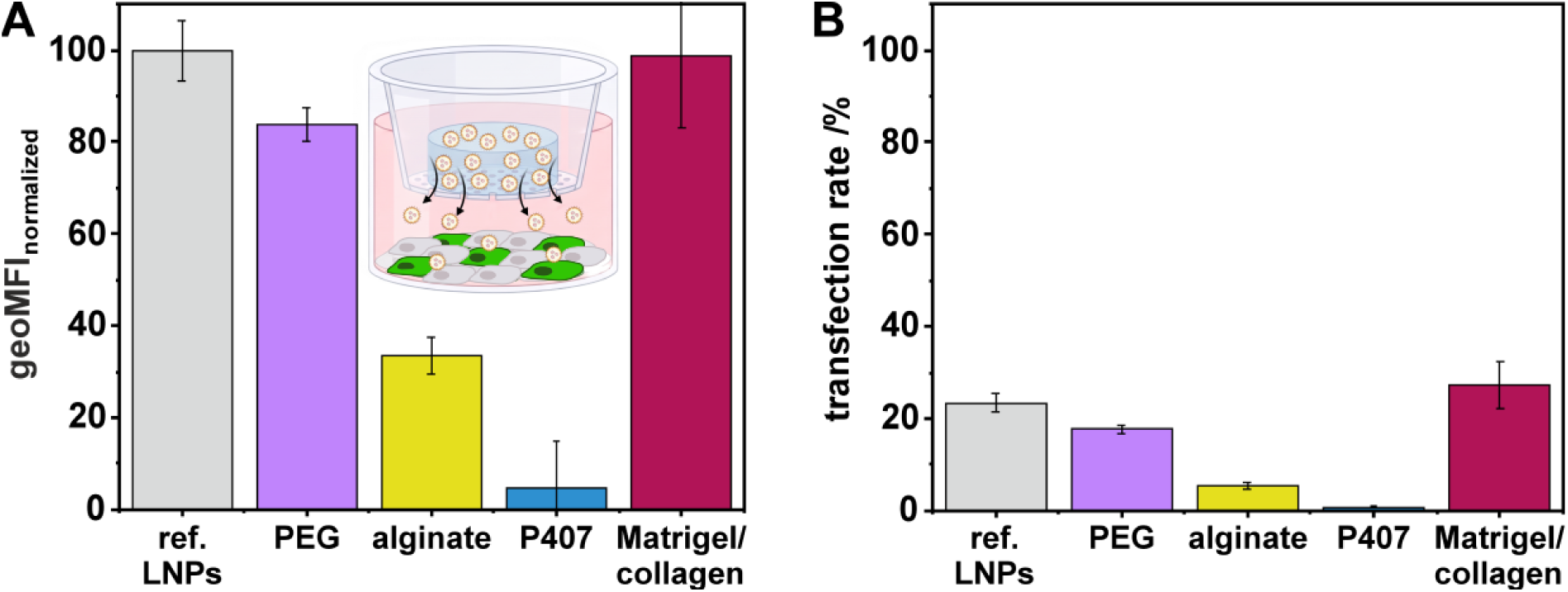
Functional transfection following continuous release of hydrogel-embedded mRNA-LNPs in a Transwell setup. mRNA-LNP-loaded hydrogels were cast into Transwell inserts and incubated above HeLa cells for 72 h, enabling continuous release of mRNA-LNPs and diffusion through the membrane to the underlying cell layer. Non-encapsulated mRNA-LNPs added directly to the insert compartment served as reference (gray). eGFP expression was quantified by flow cytometry. A) Geo Mean fluorescence intensity (geoMFI) was normalized to the reference mRNA-LNPs. PEG hydrogels largely preserved eGFP expression, whereas alginate and particularly P407 hydrogels reduced reporter expression. Matrigel/collagen hydrogels maintained fluorescence intensities comparable to the reference formulation. B) Transfection rates followed a similar trend, with the order reference LNPs ≈ Matrigel/collagen > PEG > alginate >> P407. Created in BioRender. Breunig, M. (2026) https://BioRender.com/2utsh51. Data are presented as mean ± SD.

Overall, the results demonstrate that hydrogel composition strongly influenced the biological activity of continuously released mRNA-LNPs. While PEG largely preserved functional mRNA delivery, alginate and particularly P407 impaired transfection performance. In contrast, Matrigel/collagen maintained transfection activity at levels comparable to non-encapsulated mRNA-LNPs under identical Transwell conditions.

## 4. Discussion

The present study demonstrates that hydrogel composition strongly influences the release, availability, and functional performance of mRNA-LNPs. Although all investigated hydrogels enabled the release of biologically active nanoparticles, substantial differences were observed between hydrogel classes, highlighting hydrogel composition and network architecture as key determinants of mRNA-LNP delivery.

DiI- and mRNA-loaded LNPs displayed comparable physicochemical properties, supporting the use of DiI-LNPs as a surrogate system for release studies. Stability experiments further confirmed that the LNP formulations remained largely unchanged throughout the experimental period, indicating that differences observed during release experiments primarily originated from hydrogel-dependent effects rather than intrinsic nanoparticle instability. The observed differences between mRNA-LNPs maintained under standard storage conditions (4 °C) and those incubated at 37 °C (reference LNPs and hydrogel-embedded LNPs) point towards a temperature-driven structural reorganization of the nanoparticles rather than hydrogel-induced destabilization. Increased electron density together with reduced particle diameters are consistent with a more compact internal particle structure. Such behavior may reflect relaxation of the initially formed LNP architecture from a kinetically trapped state toward a thermodynamically more favorable and denser organization at physiological temperature [41]. In this context, reorganization of the lipid components into a more tightly packed phase [41,42], potentially accompanied by core dehydration and the release of trapped water from structural defects [43], could contribute to the observed particle compaction. Importantly, fluorescence recovery measurements and fluorescence NTA yielded complementary information regarding LNP behavior after hydrogel encapsulation. While recovery of the fluorescent dye DiI was below 100 % for several formulations, NTA analysis demonstrated high particle recoveries across all investigated hydrogels. These findings suggest that reduced fluorescence recoveries did not necessarily reflect LNP loss but may partly result from leakage of DiI from otherwise intact particles. Because DiI exhibits negligible fluorescence in aqueous environments [44], fluorescence recovery can be regarded as an indirect measure of cargo retention during hydrogel release. However, DiI leakage does not necessarily imply loss of the encapsulated mRNA cargo, which is substantially larger and therefore expected to escape from the lipid structure less readily than the dye. A central objective of this work was to determine how different hydrogel microenvironments affect LNP transport and release. Although all investigated hydrogels released LNPs, substantial differences were observed between hydrogel systems, indicating that hydrogel architecture is a key determinant of LNP behavior. Across all hydrogel systems, lower polymer concentrations accelerated nanoparticle release, consistent with the established relationship between polymer content, network density, and diffusional resistance [20,45]. Importantly, these differences cannot be explained by crosslinking chemistry alone. Nanoparticle transport within hydrogels is governed by the combined effects of network architecture, degradation behavior, matrix hydration, and physicochemical interactions between the hydrogel and the encapsulated particles [46–48]. Therefore, the observed release profiles should be viewed as emergent properties of the entire hydrogel microenvironment rather than direct consequences of the crosslinking mechanism itself. Functional transfection experiments confirmed that mRNA-LNPs remained biologically active following hydrogel release. Both the degradation supernatant transfection studies and the continuous-release Transwell experiments resulted in successful eGFP expression in HeLa cells, although pronounced differences in biological activity were observed between hydrogel formulations. Importantly, similar trends were observed in both experimental approaches, indicating that hydrogel composition strongly influenced functional mRNA delivery. Diffusion across the Transwell membrane had only a minor impact on transfection performance, supporting the conclusion that the observed differences primarily originated from hydrogel-dependent effects.

As PEG is a largely inert material with limited nonspecific interactions towards nanoparticles and biomolecules [49,50], LNP mobility is expected to be governed primarily by the physical properties of the hydrogel network. Increasing polymer concentrations likely reduced nanoparticle mobility by increasing network density and diffusional resistance, resulting in progressively slower release. Accordingly, PEG hydrogels provided the broadest control over LNP availability through simple adjustment of polymer content. Despite the high LNP recovery and only moderate changes in particle size distribution, both transfection efficiency and eGFP expression decreased following hydrogel release. In the Transwell experiments, PEG hydrogels exhibited improved transfection performance compared to the degradation supernatant transfection studies. One possible explanation is that mRNA-LNPs remained protected within the hydrogel matrix until release. This observation indicates that preservation of particle number and size alone does not necessarily translate into unchanged biological performance. Although the underlying mechanism remains unclear, subtle hydrogel-induced alterations of LNP properties such as adsorption of hydrogel degradation products may have contributed to the reduced functional mRNA delivery observed after release. Nevertheless, the improved performance observed under continuous-release conditions indicates that PEG hydrogels are capable of supporting efficient mRNA-LNP delivery when sustained nanoparticle exposure can be maintained. The minor influence of PEG hydrogels on the cell viability may be attributed to the mildly acidic conditions employed during hydrogel formation to minimize hydrolysis of maleimide groups required for covalent crosslinking [51].

Alginate hydrogels exhibited comparatively low fluorescence recovery of released LNPs which indicates significant leakage of the fluorescent dye DiI. Interestingly, complete macroscopic degradation was observed during the release experiments, while alginate hydrogels remained largely intact throughout the cell uptake study. This discrepancy may be explained by the different experimental media. While release experiments were conducted in PBS buffer, the cell culture medium contained calcium ions that can stabilize the ionically crosslinked network and thereby retard hydrogel degradation. Consequently, LNP release under cell culture conditions was slower than predicted from the release studies, providing a plausible explanation for the comparatively low particle uptake observed for this material. Beyond differences in release kinetics, fluorescence NTA revealed substantially broader particle size distributions for alginate-released LNPs than for any other hydrogel system. This may reflect transient Ca^2+^-mediated interactions between polyanionic alginate and the LNP surface during hydrogel residence. Such interactions could result in small amounts of alginate molecules remaining associated with released nanoparticles, thereby increasing particle heterogeneity as detected by NTA. Together with the low fluorescence recoveries, these findings suggest that interactions between the alginate matrix and encapsulated LNPs may affect nanoparticle integrity during release. Partial destabilization of the particles could promote leakage of DiI and potentially alter the stability of the encapsulated mRNA. Although this mechanism remains speculative, it may contribute to the reduced biological performance observed for alginate formulations. Taken together, the data indicate that alginate not only modulates nanoparticle release but may also interact more strongly with encapsulated LNPs than the other hydrogel systems investigated.

P407 formulations showed little evidence of hydrogel-induced effects on the physicochemical properties of LNPs. Particle size distributions remained largely unchanged, nanoparticle recoveries were high, and cell viability was not affected by hydrogel exposure. The release behavior of P407 likely reflects the highly dynamic and reversible nature of its physically assembled micellar network, which regulates nanoparticle availability primarily through gel dissolution rather than through strong matrix-particle interactions. Consequently, P407 appears particularly well suited as a transient depot system that modulates nanoparticle retention while largely preserving the physicochemical integrity of the encapsulated LNPs. Nevertheless, transfection efficiencies of P407-released mRNA-LNPs remained below those of the non-encapsulated reference LNPs. This observation indicates that preservation of particle size and recovery does not necessarily translate into fully preserved biological performance. Notably, transfection performance was further reduced in the Transwell experiments despite continuous exposure of the cells to mRNA-LNPs released from the hydrogels. Interestingly, strong cellular fluorescence, P407-released LNPs showed only minimal transfection, suggesting that P407 permits cell association but impairs intracellular delivery required for efficient mRNA translation. Similar observations have recently been reported for P407-containing LNP formulations, where reduced reporter expression was attributed to impaired intracellular processing [52,53]. One possible explanation is reduced endosomal escape, which may limit effective cytosolic mRNA delivery despite successful association of LNPs with cells. [53,54]. In vivo, continuous dilution and clearance of the polymer are expected to reduce local P407 concentrations, which may still explain the successful application of P407-based LNP depot systems reported in previous animal studies [55,56].

The Matrigel/collagen system represented the most biologically complex matrix investigated. Quantitative assessment of LNP release was complicated by incomplete matrix dissolution, which likely resulted in an underestimation of fluorescence recovery. Nevertheless, Matrigel/collagen hydrogels supported efficient cellular uptake and yielded the highest levels of eGFP expression in this study. Collagen has been shown to exhibit high affinity toward nanoparticle surfaces and to form stable protein coronas, which may have contributed to the efficient delivery observed here [57,58]. Furthermore, collagen-rich extracellular environments have been associated with enhanced nanoparticle uptake, suggesting that collagen may promote efficient cellular interactions with the released LNPs [57,59]. The hydrogel may therefore not only act as a release matrix but also as a source of extracellular proteins capable of modifying the nanoparticle surface prior to cellular exposure [58]. In contrast to PEG, alginate and P407 hydrogels, biological activity was largely independent of polymer concentration. Notably, formulations containing low and intermediate collagen concentrations achieved eGFP expression levels approaching those of non-encapsulated reference mRNA-LNPs, demonstrating highly efficient functional delivery following hydrogel release. This finding was further supported by the Transwell experiments, in which continuously released mRNA-LNPs maintained reporter protein expression and proportions of eGFP-positive cells comparable to those achieved with the non-encapsulated reference formulation. Furthermore, cell viability remained comparable to untreated controls, confirming the expectedly excellent biocompatibility of the protein-based matrix. Together, these findings suggest that the low apparent fluorescence recoveries primarily reflect limitations of the analytical method rather than reduced nanoparticle availability. In contrast, the biological data indicate that Matrigel/collagen hydrogels effectively preserve mRNA-LNP functionality while enabling efficient cellular delivery following release.

Future LNP formulations optimized for hydrogel encapsulation may further improve biological activity following release. Thus, the transfection efficacies observed here likely represent a rather conservative estimate of the performance achievable with hydrogel-based mRNA-LNPs. Summarized, our work establishes the feasibility of hydrogel-based mRNA-LNP depots and demonstrates, through a systematic comparison of distinct hydrogel platforms, that nanoparticle release and functionality can be modulated through hydrogel design. Importantly, functional transfection was observed not only after exposure to released nanoparticles collected from degradation supernatants but also during continuous release of hydrogel-embedded mRNA-LNPs in a Transwell cell culture setup, demonstrating that released nanoparticles remain biologically active over the course of hydrogel degradation. Although the investigated release periods were limited to several days, even short-term extension of mRNA-LNP availability may be sufficient to influence antigen exposure and early immune priming. Enhancing the stability of LNPs by e.g. varying the lipid composition may make longer release periods feasible [60,61]. The present findings provide a foundation for future studies investigating antigen expression kinetics, immune responses, and vaccine performance following administration of hydrogel-encapsulated mRNA-LNPs in vivo.

## 5. Conclusion

Hydrogel encapsulation offers a versatile strategy to modulate the release and biological availability of mRNA-LNPs. The pronounced differences observed between hydrogel classes demonstrate that matrix composition and network architecture are key determinants of nanoparticle release and functionality. Importantly, all investigated hydrogels released biologically active mRNA-LNPs capable of mediating protein expression while maintaining high cell viability, establishing the feasibility of hydrogel-based mRNA-LNP depots. We expect LNP formulations specifically tailored for hydrogel encapsulation to further enhance biological activity following release. These findings establish design principles for the development of hydrogel-based mRNA delivery systems and highlight the potential of biomaterial-assisted approaches to prolong mRNA-LNP availability.

## CRediT authorship contribution

**Andreas G. Schreiber:** Writing – review & editing, Writing – original draft, Visualization, Methodology, Investigation, Formal analysis, Data curation, Conceptualization. **Felix Hauswirth:** Methodology. **Leon Reger:** Methodology. **Simone P. Carneiro:** Methodology **Olivia M. Merkel:** Writing – review & editing, Supervision. **Miriam Breunig:** Writing – review & editing, Supervision, Investigation, Funding acquisition, Conceptualization.

## Declaration of competing interests

The authors declare that they have no known competing financial interests or personal relationships that could have appeared to influence the work reported in this paper.

## Acknowledgements

Financial support from the Deutsche Forschungsgemeinschaft (DFG, grant No. 3566/5-1) is gratefully acknowledged. The authors thank Lipoid GmbH for kindly donating DSPC and Renate Liebl as well as Grace Faltermeier for outstanding technical assistance. We acknowledge Otto Berninghausen of the Gene Center, LMU Munich, for cryoTEM image acquisition.

## Declaration of generative AI and AI-assisted technologies in the manuscript preparation process

During the preparation of this work the authors used Microsoft Copilot (GPT-5) for language editing and text refinement. After using this tool, the authors reviewed and substantially edited the content and take full responsibility for the content of the published article.

## Notes

### Competing Interest Statement

The authors have declared no competing interest.

### Summary of Updates

Additional experiments were performed to strengthen the study and provide further insight into the stability, structure, and cellular interaction of mRNA-loaded lipid nanoparticles (mRNA-LNPs) released from hydrogels. New CryoTEM analyses were included to assess potential morphological changes of mRNA-LNPs following incubation at physiological temperature and after release from hydrogels. These data revealed temperature-dependent structural changes. In addition, a new Transwell experiment was performed to assess the functional delivery of hydrogel-released mRNA-LNPs. The observed GFP expression demonstrated that mRNA-LNPs released from hydrogels remained biologically active and retained their capacity to mediate protein expression following hydrogel-mediated release.

