## Supplementary information for "Hydrogel crosslinking mechanisms influence the release and functional delivery of lipid nanoparticles"

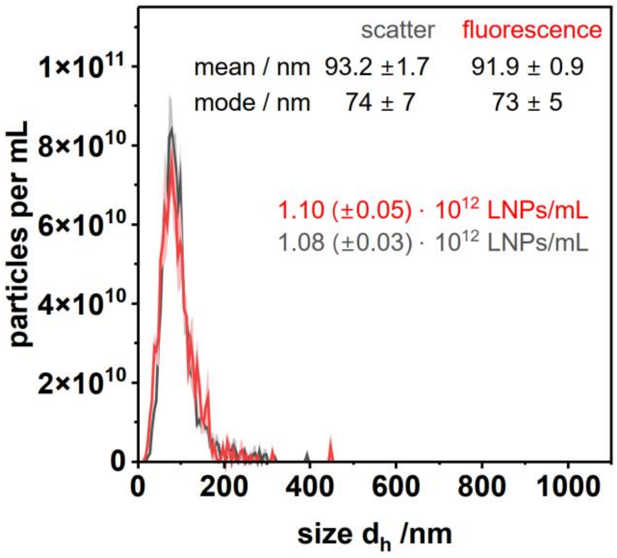


**Fig. S1 Comparison of light scattering and fluorescence NTA analysis of DiI-LNPs.** Particle size distribution and particle concentration determined by nanoparticle tracking analysis were comparable in light scattering and fluorescence detection modes.

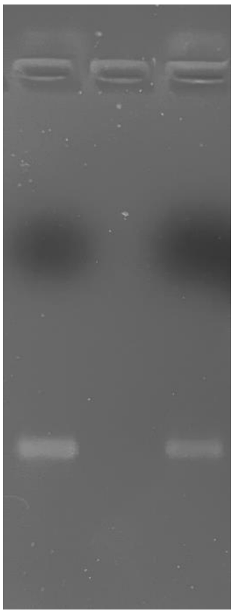

**Fig. S2 Agarose gel electrophoresis of mRNA.** A 1 % agarose gel was used to compare fresh mRNA (lane 1, left) with mRNA recovered from mRNA-LNPs following particle lysis with Triton X 100 (lane 2, right). Comparable migration patterns were observed for both samples.

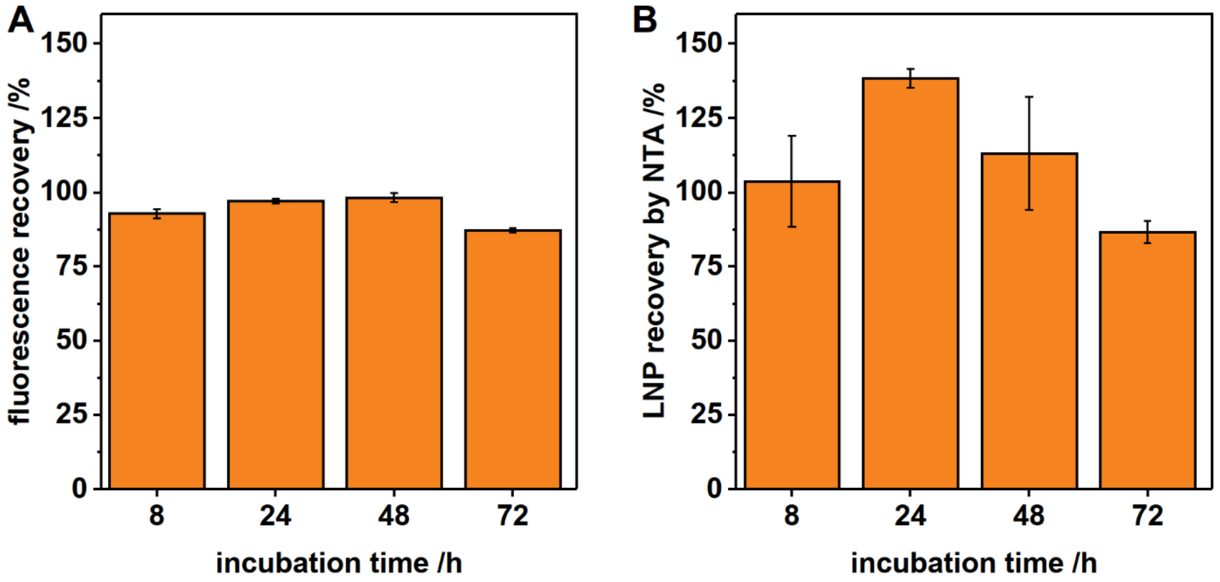


**Fig. S3 Stability assessment of DiI-LNPs under release conditions.** DiI-LNPs were diluted in PBS as uses for in the release experiments (15 µL LNP suspension adjusted to a total volume of 400 µL) and incubated at 37 °C. A) Fluorescence intensity of DiI-LNPs remained constant over 72 hours. B) Particle concentration determined by fluorescence NTA showed no significant changes over time. Data are presented as mean ± SD.

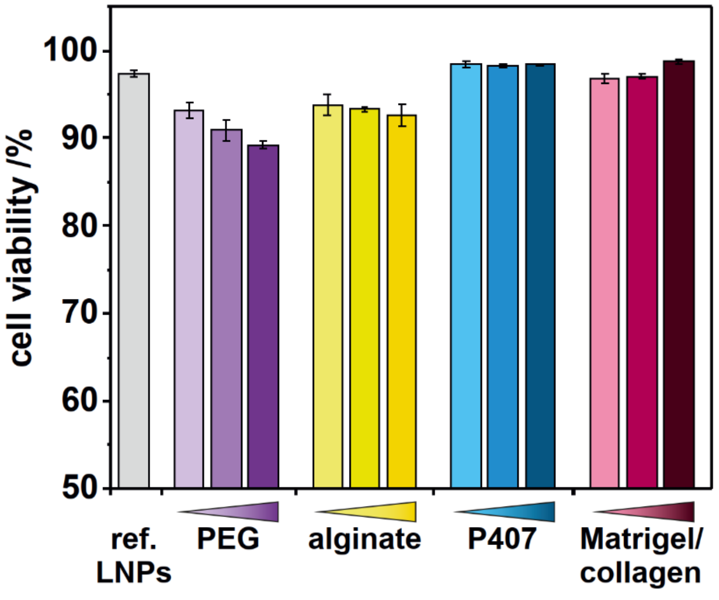


**Fig. S4 Assessment of hydrogel-associated cytotoxicity by flow cytometry.** The supernatants of hydrogels without LNPs after degradation for 72 h at 37 °C were applied to HeLa cells. After 24 h of incubation, cell viability was determined by propidium iodide staining and flow cytometry. PEG and alginate hydrogels exhibited a slight reduction in cell viability, whereas all other hydrogel formulations maintained viabilities comparable to the PBS control. Data are presented as mean ± SD.

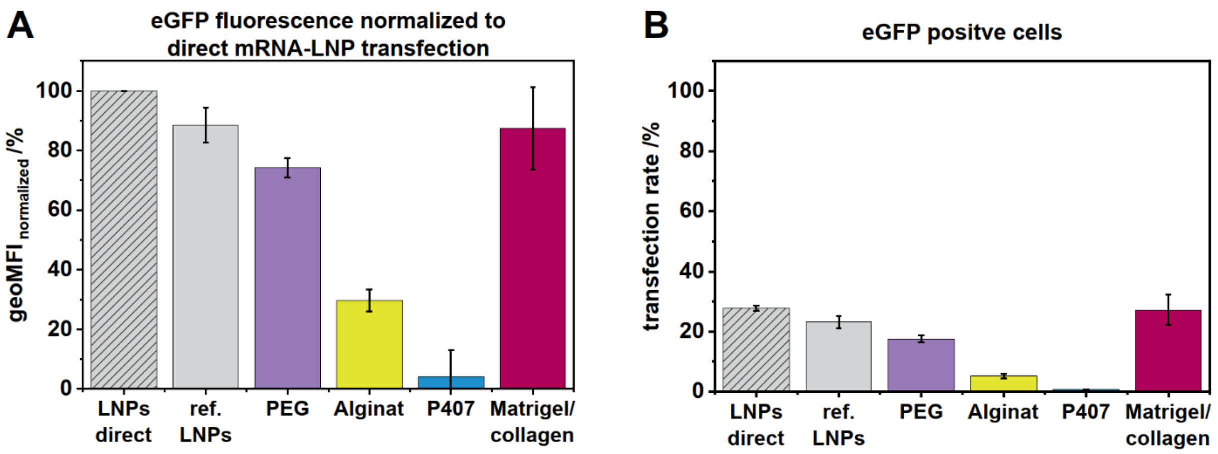


**Fig. S5 Comparison of direct and Transwell-mediated mRNA-LNP delivery**. Non-encapsulated mRNA-LNPs were either added directly to HeLa cells (striped gray) or to the insert compartment of a Transwell system (gray). mRNA-LNP-loaded hydrogels were cast into Transwell inserts. All samples were incubated above HeLa cells for 72 h, enabling continuous release of mRNA-LNPs and diffusion through the membrane to the underlying cell layer. eGFP expression was quantified by flow cytometry. A) Geo Mean fluorescence intensity (geoMFI) was normalized to directly applied mRNA-LNPs (striped gray), which were set to 100 %. Transwell reference mRNA-LNPs exhibited eGFP expression levels comparable to directly applied mRNA-LNPs, indicating that passage across the Transwell membrane had only a minor impact on reporter expression. B) Transfection rates followed a similar trend. Direct mRNA-LNPs and Transwell reference mRNA-LNPs resulted in comparable percentages of eGFP-positive cells, confirming the observations made for normalized eGFP expression in panel A and indicating that passage across the Transwell membrane had only a minor effect on transfection efficiency. Data are presented as mean ± SD.
